# Predictable sequential structure augments auditory sensitivity at threshold

**DOI:** 10.1101/2023.10.03.560670

**Authors:** Nadège Marin, Grégory Gérenton, Hadrien Jean, Nihaad Paraouty, Nicolas Wallaert, Diane S. Lazard, Keith B. Doelling, Luc H. Arnal

## Abstract

Human hearing is highly sensitive and allows us to detect acoustic events at low levels. However, sensitivity is not only a function of the integrity of cochlear transduction mechanisms, but also constrained by central processes such as attention and expectation. While the effects of distraction and attentional orienting are generally acknowledged, the extent to which probabilistic expectations influence sensitivity at threshold is not clear. Classical audiometric tests, commonly used to assess hearing thresholds, do not distinguish between bottom-up sensitivity and top-down processes. In this study, we aim to decipher the influence of various types of expectations on hearing thresholds and how this information can be used to improve the assessment of hearing sensitivity. Our results raise important questions regarding the conventional assessment of hearing thresholds, both in fundamental research and in audiological clinical assessment.

## Introduction

The sense of hearing allows us to detect and interpret even very weak sounds in our environment. Typically, sensitivity is primarily attributed to the intricate mechanics of the middle and inner ear, and the synaptic transduction of signals to the auditory nerve (1–3). The efficacy of these mechanics is largely measured using pure-tone detection (PTD) thresholds, task paradigms designed to gauge the softest intensities at which a person can detect pure tones, which are standard practice in clinical audiology (4).

In recent decades, neuroscience has increasingly embraced the idea that sensory processing is more than just a result of bottom-up signal transduction. According to predictive processing theories (Friston, 2005), perception is an active process heavily relying on our capacity to anticipate upcoming events based on our sensory experience. These predictions are proposed to be organized in a hierarchical manner across various levels of abstraction, such as acoustic, phonemic, or semantic levels, for instance (5, 6).Top-down signals proactively filter and modulate the gain of expected stimulus inputs (7). As such, even without conscious awareness, the human brain extracts statistical regularities to exploit predictable structures in a task (8). Does the effect of prediction extend to even the most fundamental and low-level auditory function, pure-tone sound detection?

Previous work has shown that hearing sensitivity is contingent on more than the mere transduction of ascending signals at the cochlear level but is also affected by the allocation of cognitive constructs such as attentional resources towards upcoming events depending on their behavioural relevance. The probe-signal task (9), for example, demonstrated that detection of a tone is enhanced when it is preceded by a probe tone around the same frequency. A wealth of studies in psychoacoustical experimentation have demonstrated this improvement in performance based on this sort of selective attention in noise (10–12). These studies operate under the assumption that the effects of top-down modulation of signal detection would be the same in quiet at threshold and that the added noise merely externalizes and enhances the neural noise of the system.

On the other hand, the audiological field considers the pure-tone audiogram (signal detection in quiet) to diagnose only the bottom-up sensory pathway (13), even though current audiometry protocols contain considerable predictive structure. Do the findings of psychoacoustics in noisy settings apply to signal detection in quiet? Or does expectation only play a role when separating auditory signal from acoustic noise?

While previous work has studied the role of attention in quiet, this has largely been in terms of attention to hearing vs visual input. Rather than manipulating the expected information of attended stimuli, these studies consider attention as a means of removing distraction from one or another sensory domain. Lukas (14), for example, demonstrated that brainstem responses from auditory nerve to colliculus were filtered by sustained concentrated attention to the visual domain, suggesting top-down control to avoid distraction. Later evidence further suggested that attention has cochlear effects, demonstrated by changes in oto-acoustic emissions during attention to one ear vs the other (15) and by changes in rhythmic modulation of the auditory nerve fiber during attention to the auditory vs visual domain (16). These findings provide a mechanism for top-down modulation of cochlear sensitivity via the olivocochlear bundle. Given that a decidedly cortical construct can have direct effect on peripheral function, it is reasonable to explore whether other more stimulus-specific cognitive functions may also influence cochlear processing.

Of particular importance in the case of audition is the perception of not only the content of a sound but also its timing. For example, speech contains prosodical cues such as the speeding up or slowing down of syllabic rate which provides further nonverbal information to the listener. As such, predictions about an upcoming sensory event can not only regard its content but also its moment in time. This delineation between content-based (“what”) and time-based (“when”) predictions has been maintained in the field suggesting that the two rely on different top-down mechanisms (17). In the case of timing, a growing body of evidence supports the coupling of neural oscillations in different timescales to statistical regularities in the sensory input as a mechanism for predictive timing, particularly in the case of auditory stimuli (18–20). Meanwhile, content predictions are sent as inhibitory top-down signals (21–23), reducing the processing of predicted signals so that only what is unexpected is given greater neural resources.

These components of prediction, while developed over a large body of research across several decades in the fields of psychology and neuroscience, have not been heavily considered in the audiological domain. In the clinic, one baseline measurement for auditory health focuses on pure tone detection, identifying the softest presentation level at which a participant can detect a pure tone. However, such measurements overlook the predictive nature of perception, often presenting the same tone repeatedly and regularly, thereby allowing for the participant to predict when and what a tone is before it arrives. Recent protocols in audiological paradigms have begun to take this into account, instructing clinicians to randomize the timing of tones in pure tone detection tasks to avoid guessing (24). Still, content predictions (i.e., pitch) are ignored and the role such predictions play and how they interact with temporal predictions in threshold evaluation is unknown. While much work on prediction has focused on signal recovery in noisy environments, very little work has been done at auditory detection threshold in quiet. Our goal is to assess how much sequential structures contribute to thresholds evaluation under current clinical techniques and to what extent the predictions in either time or content can affect sensory outcomes.

In this study, we take advantage of new advances in pure tone detection paradigms (25) by employing automated detection threshold techniques using Bayesian Machine Learning to infer audiograms based on a more flexible paradigm, allowing for a fully randomized structure in both timing and frequency. We use this advance to compare audiograms within subject, conducting two experiments in which participants undergo audiometric testing. First, we compare directly a gold standard clinical and psychophysical paradigm and the fully randomized paradigm we developed. Then, in a second experiment we compare the effect of predictability more directly by contrasting predictability in time and frequency in a highly controlled format at slow (1-3 s) and fast temporal scales (400-800 ms).

In contrast with current clinical expectations, we expect that predictive structure will facilitate sensory detection at threshold. This experiment will provide key information regarding the role of prediction in detection at auditory thresholds in the normal hearing listener and will therefore provide the foundation for future work to show how this relationship may be altered in hearing-impaired populations.

## Methods

### Participants

31 participants aged 18 to 45 with self-reported normal hearing were recruited using the RISC (Relai d’Information sur les Sciences de la Cognition) volunteers’ database (Risc., https://www.risc.cnrs.fr/.). Normal hearing was confirmed (Pure Tone Average < 20 dB HL) in the first audiometry test (described below) and no participants were removed on this basis. Three participants were excluded from all the analyses, for either not following the instructions, or due to a data collection error. We present the results from the remaining 28 participants (12 men, 16 women) with an average age of 26.25 years-old (sd: 5.73). Informed consent was obtained prior to testing and participation in this study was compensated. One session lasted about 1h30 in total. We assessed self-reported musical ability by administering the Goldsmiths Musical Sophistication Index (Gold-MSI). This study was approved by the Comité de Protection des Personnes Tours OUEST 1 on 10/09/2020 (project identification number 2020T317 RIPH3 HPS).

### Stimuli and Material

All experiments were created using the open-source python library Psychopy 2022.1.1 (26). Stimuli consisted of 200 ms pure tones with frequencies spanning the 125-8000 Hz range. Stimuli were presented binaurally in a sound attenuated booth, through Etymotic Research ER-2 insert earphones fitted with 13mm Echodia plugs and connected to a Solid State Logic SSL 2 soundcard. The 200ms pure tones were apodized with 5ms hanning windows and hearing thresholds were measured for tones calibrated to ISO 389-2, with reference to an IEC-711 occluded-ear simulator. Psychopy’s sound module interprets loudness as a number comprised between 0 and 1, therefore the calibration was performed for an arbitrary Psychopy volume of 0.005 before applying the correction to dB HL. The presentation level was linearly interpolated for frequencies not included in the correction table.

### Experimental setup

In experiment 1, participants completed two audiometrical tasks : a Randomized paradigm and a 3-alternative forced choice (3-AFC) adaptive staircase paradigm. The 3-AFC adaptive staircase (Figure 1A, and detailed below) was designed to reproduce the current gold-standard in audiometry procedures and allow us to compare its threshold estimation with other assessments made using unpredictable tones. The Randomized task (Figure 1B) measures thresholds for pure tones as unpredictable in timing and frequency as possible within this study’s limitations.

**Figure 1-.**
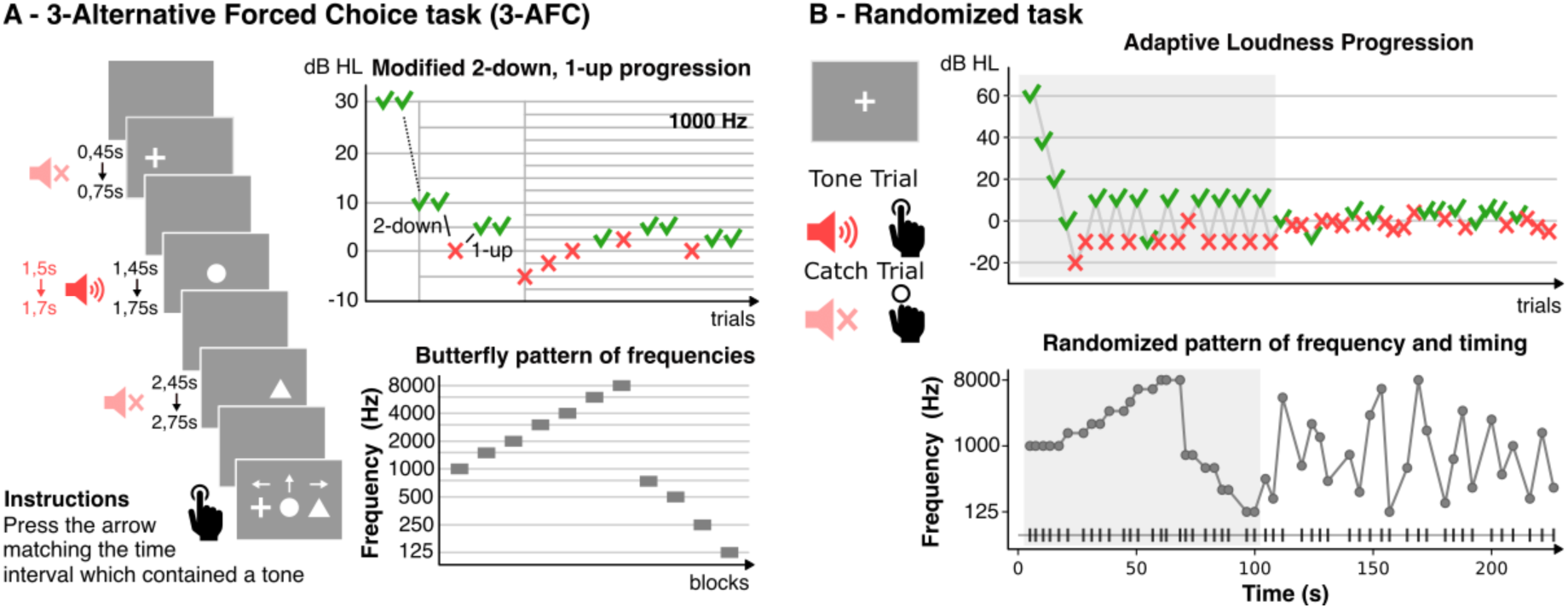
Experiment 1: predictable and unpredictable designs. ***A – 3-AFC task.*** *Participants were asked to identify which of 3 possible time intervals contained a tone. A 2-down, 1-up adaptive staircase starting at 30 dB HL determined the level of the presentation of the tones. There is structure in the order of presentation of frequencies, as the next frequency is chosen following a butterfly pattern. The design of this task carries a lot of predictability, both in time and in content. **B – Randomized task.** A fixation cross was displayed on screen while tones chosen by an automated pure-tone audiometry procedure were presented with randomized ISI (1-3 sec). Instructions were to press a key as soon as a tone was detected. In this task, the timing and frequency of successive tones is unpredictable*.

In experiment 2, the same participants completed two additional audiometrical tasks: a fast-paced paradigm, and a slow-paced paradigm, which recorded 4 audiograms each, in different conditions of predictability in time and frequency.

As all volunteers participated in both experiment 1 and 2, we used the randomized paradigm in experiment 1 to confirm inclusion in the study (< 20 dB HL hearing loss) and to choose the sound presentation levels in both paradigms of Experiment 2 (see below). To estimate thresholds, all tasks except for the 3-AFC adaptive staircase of Experiment 1 rely on a state-of-the-art, automated pure-tone audiometry procedure by My Medical Assistant SAS (iAudiogram, www.iaudiogram.com) validated against customary adaptive staircase procedures and based on the automated machine learning described previously (25).

#### Experiment 1: Comparison of Audiometrical Threshold Estimation Paradigms

##### Three Alternative Forced Choice (3-AFC) paradigm

At each trial, participants were asked to identify one of three possible time intervals during which a tone had been played. A sequence of 3 shapes flashed on screen (a cross, a disk and a triangle respectively on the left, in the middle and on the right) to indicate the three intervals in time (Fig. 1A, left). Each shape was displayed during 300 ms and the next one appeared after a delay of 700 ms. Participants had to press the keyboard arrow corresponding to the correct interval, randomly chosen on a trial-by-trial basis. The experiment started with five practice trials to familiarize participants with the task. We tested 11 frequencies ordered in a butterfly pattern spanning the 125-8000 Hz range (1000 Hz, 1500 Hz, 2000 Hz, 3000 Hz, 4000 Hz, 6000 Hz, 8000 Hz, 750 Hz, 500 Hz, 250 Hz, 125 Hz) (Fig. 1A, bottom right). The first tone of each frequency was presented at 30 dB HL. The level of presentation of the next tone was determined based on a 2 down – 1 up adaptive staircase procedure (Fig. 1B, top right). The initial step of 10 dB HL was reduced to 5 dB HL after the first decrease in level. We halved the step again after 3 reversals of the staircase and stopped after 6 reversals. Participants had the opportunity to take a short break before starting the next frequency’s staircase. Pure tone thresholds were estimated as the average level across the last 4 reversals. The task lasted on average 20.11 minutes (sd: 3.36 minutes).

##### Randomized paradigm

Participants were asked to detect a tone at low decibel levels, indicating their detection by pressing a key when they heard the tone (Fig. 1B, top). They were informed that detections would only be considered correct if they occurred within a 1-second window following the tone. A fixation cross was displayed on the screen during the whole task. Participants were familiarized with the paradigm using three practice trials at 40 dB HL. The task starts with an initialization phase testing tones spanning the frequency range (1000Hz, 1500Hz, 2000Hz, 3000Hz, 4000Hz, 6000Hz, 8000Hz, 750Hz, 500Hz, 250Hz, 125Hz) to provide an initial coarse estimation of the audiogram. During this phase, the frequency of tones is predictable, as one frequency is tested several times in a row at decreasing hearing levels, until a negative answer is recorded for that frequency. After the initialization phase, the following sound presentation levels and frequencies were chosen to maximally reduce the uncertainty of the threshold estimation using Bayesian inference (Fig. 1B, bottom). The estimation is based on recent work designed to automate pure tone detection threshold (25). It uses a Gaussian Process with prior knowledge about the relationship between points in the frequency-level space, to assign a probability that a tone of any level and frequency will be heard. The prior knowledge is encoded through kernels: one with a covariation matrix using a squared exponential in frequency space, denoting that neighboring frequencies are correlated, and another as a linear function in intensity space, denoting that increased loudness should increase the probability of detection. The next tone is chosen as the point leading to the greatest decrease in the uncertainty of these probabilities, so its frequency cannot be predicted by the participant. The timing in between each tone, the Inter-Stimulus Interval (ISI), was randomly chosen for each tone to be between 1 and 6 seconds to reduce the participants’ ability to predict the timing of upcoming tones. Each tone had a 20% chance of being replaced with silence to be analyzed as catch trials, a fact that participants were not informed about. The true average percentage of catch trials across participants amounted to 18.81% ± 1.57% of trials. The paradigm completes when the Bayesian estimator reaches a confidence interval of 6 dB. With this constraint, the complete paradigm comprised an average of 60.9 trials (min: 51, max: 101, sd: 17.9), including an initialization phase of 25.1 trials (min: 22, max: 28, sd: 1.4), for a total average duration of 6.24 minutes (sd: 2.20 minutes).

##### Analysis

The 3-AFC and Randomized paradigms use different algorithms to calculate the final threshold: Adaptive Staircase and Bayesian Active Learning, respectively. These methods differ in the continuity of the outputted thresholds (the staircase is necessarily discrete, whereas the Bayesian algorithm is continuous), but otherwise are readily comparable. As such, to account for this difference we sample the threshold values of the Randomized paradigm at the frequencies tested in the 3-AFC. We then average the values across frequency for each participant to get a mean detection threshold comparable with the 3-AFC in two ways: first, averaging across all 11 frequencies of the 3-AFC, and second, averaging over the 4 critical frequencies commonly tested to estimate the pure tone average (PTA) in the clinic, 500, 1000, 2000 and 4000 Hz. The difference in thresholds from each metric are compared using a T-test.

While the 3-AFC is designed to infer a threshold at 70.7% correct, this percentage corresponds to a 56.5% chance of tone detection (accounting for guessing at 33% when tones go undetected). We consider this percentage to be negligibly different from the 50% chance inferred by the Randomized. Furthermore, correcting for the 6.5% difference, if possible, would only enhance the size of the effect between the two conditions as we expect the more predictable task (3-AFC) to result in lower thresholds.

#### Experiment 2: Effects of Temporal and Frequency Predictability on Audiometrical Detection

For both the slow- and fast-paced paradigms in experiment 2, tones were organized in ‘sweeps’ whose structures defined four conditions of predictability: predictable timing (T), predictable frequency (F), predictable frequency and timing (FT), or random frequency and timing (R) (Fig. 2A, B). The main difference between these two tasks is the timescales (ISI) used to separate successive tones. In the slow-paced task, we recorded whether participants detected each tone within a sweep, to emulate clinical procedures where an answer is given for each tone. This required using large ISIs to allow time for participants to answer. Each tone thus consisted in a trial and each sweep represented one block. In the fast-paced task, meanwhile, each trial tested the detection of a target tone preceded by a rapid cluster of 4 cue tones, in a fashion more relevant to prediction studies that tend to use shorter ISIs. In this task, one sweep represented one trial.

**Figure 2-.**
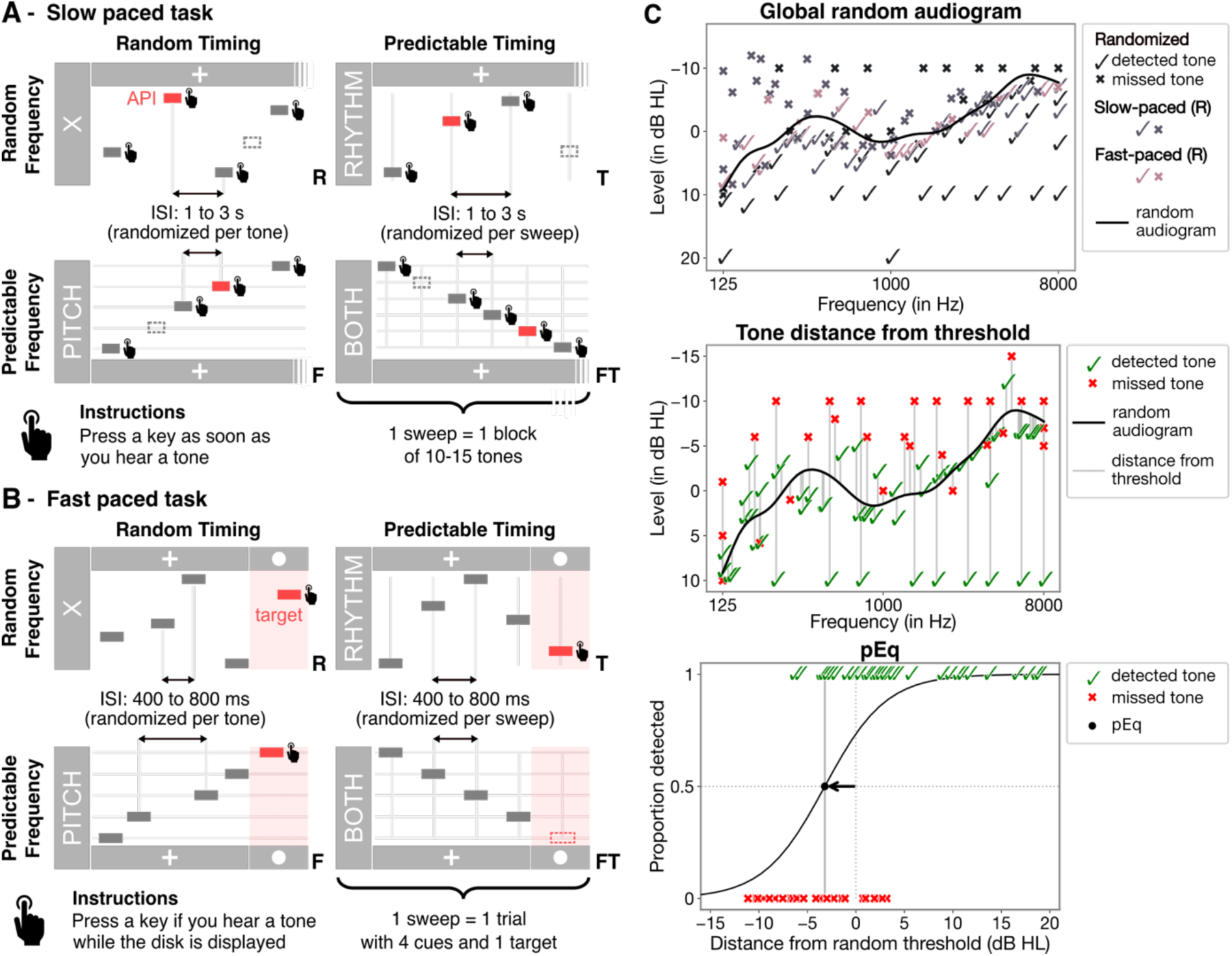
Experiment 2: Slow- and fast-paced designs. *Four conditions of predictability were established for both paradigms: predictable timing (T), predictable frequency (F), predictable frequency and timing (FT), or random frequency and timing (R). Predictability in each modality was achieved by designing sweeps with constant intervals, i.e., by regularly spacing tones in time and/or frequency. In conditions of unpredictability, we randomized the intervals (ISIs and/or frequency intervals) for each tone in a sweep. **A – Slow-paced task.** Sweeps consisting of 8 to 15 tones were presented while a fixation cross was displayed on screen. The instructions were to press a key as soon as a tone was detected. Each sweep contained one catch trial. **B – Fast-paced task.** A cluster of 4 cue tones served to indicate the timing and frequency of a 5th target tone, or not. A fixation cross was displayed on screen until the end of the last cue. A disk then replaced the cross to visually indicate the interval during which the target tone could be presented. Participants were to press a key if they detected a tone while a disk was displayed on screen. 20% of trials contained no target tone, to serve as catch trials. **C – Method for estimating pEq values (example data from one participant).** Top panel: A ‘global random’ audiogram (solid black line) is computed for each participant from aggregating the answers of the Randomized task and condition R of the Slow- and Fast-paced tasks. Middle panel: for each tested tone in a sweeping condition (R, F, T, or FT; checkmarks: positive detections, crosses: missed detections), the distance from the ‘global random’ threshold at the tested tone’s frequency within each condition is estimated in dB HL (vertical grey lines). Bottom panel: The tones tested in each condition are ordered by their distance from the ‘global’ threshold and their proportion of detection is estimated by fitting a sigmoid to the participant’s answers. We defined pEq as the distance from the ‘global random’ threshold corresponding to the 50% mark of detection*.

The four conditions were obtained by building sweeps using the same structure in both paradigms: we made sweeps predictable by regularly spacing their tones in time and/or frequency. For predictable timing, the ISI of a sweep was kept constant and was randomly selected from a paradigm-dependent range (Slow-paced: 1-3s, Fast-paced: 400-800ms). For frequency predictability, the distance between two successive tones was picked at random from a list of three musical intervals: (whole tone, major third or perfect fifth) and was the same for all tones in a sweep. To create unpredictable sweeps, the intervals between successive tones were randomized (ISIs for time and musical intervals for frequency) from the same distributions chosen for predictable sweeps. We subsequently shuffled the order of the tones for sweeps with unpredictable frequency (T, R).

We measured a separate audiogram in each of the four conditions of predictability and the order of presentation of the conditions was randomized for each sweep. Participants were made aware of the predictability of each sweep through a visual cue. The word ‘PITCH’ would flash on screen for 1 second before sweeps with predictable frequency, and the word ‘RHYTHM’ flashed the same way for sweeps with predictable timing. Both words flashed simultaneously before FT sweeps, while fully random sweeps were preceded by a flashing cross.

##### Further details on slow-paced paradigm

Instructions were the same as those for the Randomized task: participants had to press the spacebar less than 1 second after the onset of tones they managed to detect. Stimuli were presented in sweeps of 8 to 15 tones organized in time and pitch, or not (Fig. 2A). After flashing the next sweep’s condition for 1 second, a fixation cross was displayed until the end of the sweep. Each sweep was centered around a key tone, positioned near the middle in predictable frequency conditions and chosen by the same algorithm used in the randomized task to ensure coverage of tones where the model was most uncertain. Initially, sweeps were designed with 15 tones, with the key tone in the 8th position. If tones fell outside the 125-8000 Hz range, sweeps were truncated and redesigned to maintain at least 8 tones. The presentation levels of non-key tones were randomly set between their detection threshold (established in the prior randomized task from Experiment 1) and the level of the key tone chosen by the model based on data from this paradigm. ISIs were randomly chosen from a range of 1 to 3 seconds. Each sweep contained one catch trial which replaced a randomly chosen tone in the sweep from the third position onward (9.20% ± 0.99% of tones across participants). Data from the first 2 tones of each sweep were excluded from analyses in the F, T, and FT conditions since they cannot be predicted using previous tones. The average duration of this paradigm was 15.52 minutes (sd: 3.55 minutes).

##### Fast-paced paradigm

Participants were instructed to report whether they could hear a tone while a disk was displayed on the screen. Before the disk was displayed, a cluster of four pure tones presented at the same time as a fixation cross served as cues for the target tone. For trials with random timing (F, R), the delay after each of the four cues was drawn from a uniform distribution between 400 and 800 ms. In predictable-timing conditions (T, FT), the delay was consistent per trial (per sweep), randomly selected on each trial to be between the same 400-800 ms range. The disk appeared immediately after the last cue ended and remained displayed during 1.2s. Only key presses collected within one second of the target tone’s onset were considered positive detections. Participants were familiarized with the experiment in an initial training phase consisting of five mock trials with feedback.

We again used the algorithm from the randomized task in Experiment 1 to choose the presentation level and frequency of the target tone in every other trial. The target frequency of the remaining 50% of trials was randomly picked from a log-uniform distribution ranging 125 to 8000 Hz, while the hearing level was interpolated from the audiogram estimation made in the Randomized paradigm to ensure that selected tones could not be predicted by any unrealized patterns in Bayesian Inference model. The level of the cues was also interpolated from the Randomized audiogram and raised by 6 dB to increase the probability of their detection. Each target tone had a 20% chance of being replaced with silence to be analyzed as a catch trial (19.53% ± 6.47% of trials across participants). This paradigm lasted an average of 25.01 minutes (sd: 8.91 minutes).

##### Analysis

###### Point of Equivalence estimation

After the initial comparison in experiment 1, it is important to control for the difference in methods used to assess thresholds and compare all paradigms under the same footing. As such, when considering all paradigms together (Randomized, 3-AFC, and slow and fast-paced paradigms), we first calculate a global threshold using Bayesian Active Learning as in the Randomized paradigm incorporating tones from all random conditions (Randomized paradigm of Experiment 1 and Random conditions from Experiment 2 protocols - Figure 2C, top panel). This threshold reflects an audiogram from unpredictable stimuli without dependence on the paradigm protocol. Then for each condition of the slow- and fast-paced tasks (FT, F, T and R), we consider the distance of each tone presented in terms of its intensity from the global threshold (Figure 2C, middle panel). We use this distance as an independent variable in a logistic function to predict whether each tone will be detected or not (Figure 2C, bottom-panel). If the point of equivalence (pEq) of the logistic function is equal to 0, then there is no difference in threshold for this condition relative to global threshold. However, if there is a significant difference in either direction, we can assume the threshold has shifted by this amount. This method allows us to treat each condition in the same manner comparing its outcome as a relative distance from all conditions with unpredictable stimuli.

###### Statistics for Experiments 1 & 2

All statistical analyses were conducted using Python libraries, including SciPy (27), Statsmodels (28), Pingouin (29) and Scikit-learn (30). In experiment 1, audiograms were compared across frequencies and paradigms using two-factor repeated-measures ANOVA, with Greenhouse-Geisser correction applied where the sphericity assumption was violated. Paired t-tests were performed to compare mean thresholds across frequencies and paradigms, using the Benjamini Hochberg method to correct for False Discovery Rate (FDR) (Benjamini & Hochberg, 1995).

To ensure a consistent comparison across frequencies, paradigms and conditions, all subsequent analyses in experiment 2 made use of the calculated pEq values, which serve as a normalized metric representing the relative deviation of the detection threshold from the reference threshold. Separate one-way ANOVAs were implemented for the fast- and slow-paced paradigms to assess differences within paradigms based on predictability levels. Post-hoc pairwise comparisons with Benjamini Hochberg correction were performed to determine significant differences between different conditions.

To investigate potential decisional bias, we incorporated catch trials into all paradigms except the 3-AFC task and computed false alarm rates from these trials. We conducted a repeated-measures ANOVA, treating each testing condition – be it the single condition in the Randomized task or any of the four distinct conditions in the fast- and slow-paced paradigms – as a separate group. To identify significant variations in false alarm rates across these conditions, we performed post-hoc comparisons using False Discovery Rate correction (31). We performed Pearson’s correlation analysis to investigate relationships between false alarm rates and pEq threshold differences and between participants’ characteristic features (age and musicianship) and their ability to benefit from predictive information. In all tests, the significance level was set at p < 0.05. We also used unsupervised hierarchical agglomerative clustering to evaluate the patterns of participant performance across paradigms and predictability conditions. Clustering was performed using SciPy’s hierarchical clustering module with linkage using an average method and cosine distance to avoid clustering based on overall performance. All data are presented as means ± standard error of the mean.

## Results

### Experiment 1

#### Predictability augments audiometric thresholds by 7 dB

First, we compared audiograms from the Randomized paradigm (estimated using Bayesian Machine Learning) and the predictive, 3-AFC paradigm (two-down, one-up adaptive staircase), which represent our most extreme cases of random and predictable respectively, while also being the most standard in terms of previous testing. Figure 3A shows the average audiogram of all participants for the two paradigms across the tested frequency range from 125 – 8000 Hz. The result shows a significant difference in intensity across frequencies in the two paradigms (two-factor repeated-measures ANOVA, paradigm effect: F(1, 27) = 271.50, p < 0.001; frequency effect: F(10, 270) = 11.14, p < 0.001; paradigm x frequency interaction: F(10, 270) = 2.44, p = 0.009). To estimate the size of this effect, we then extracted, from the Randomized threshold, the discrete threshold values at frequencies that were also tested in the 3AFC and compared the mean values. Figure 3B shows that this comparison yielded a mean improvement of 6.4 dB HL (t(27) = −13.24, p < 0.001) for the 3AFC paradigm. We repeated this analysis using only the thresholds measured for the frequencies 500 Hz, 1000 Hz, 2000 Hz and 4000 Hz to compute the Pure Tone Average (PTA), a commonly used metric in audiology. For these selected frequencies, the mean improvement in thresholds from Randomized method to the 3-AFC was 7.19 dB HL (t(27) = −14.41, p < 0.001).

**Figure 3-.**
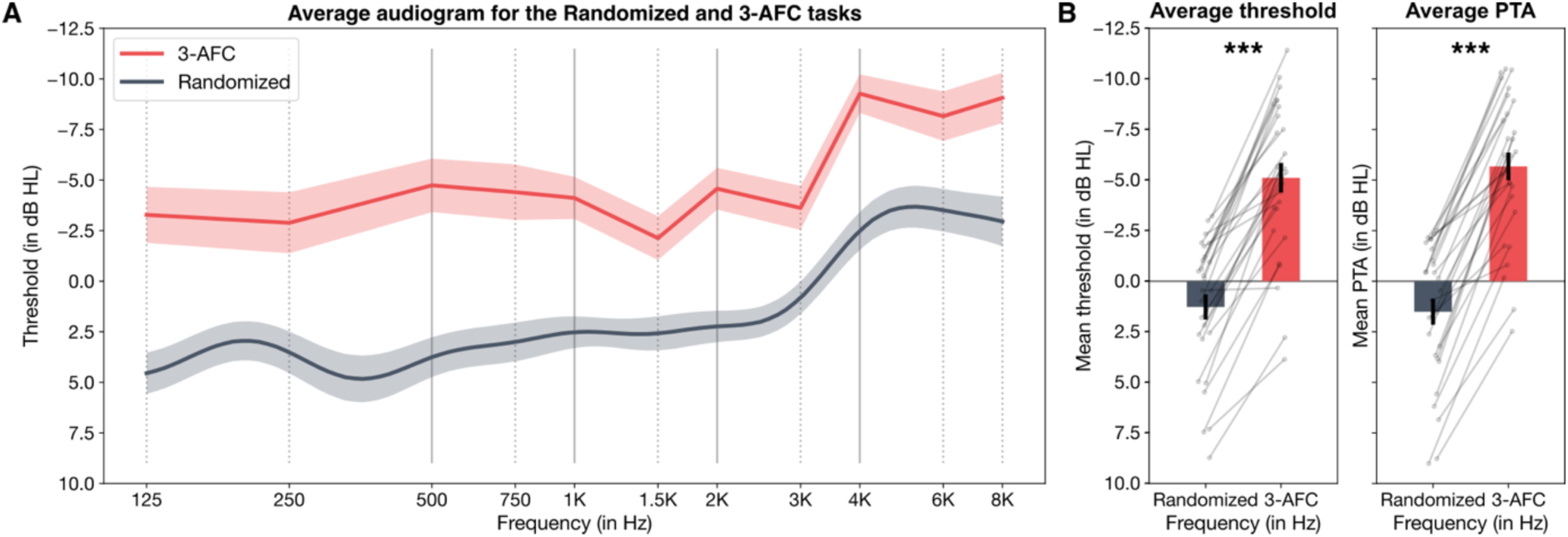
Experiment 1: Thresholds are lower in the highly structured 3-AFC task than in the Randomized task. *A – Average audiograms measured in the Randomized task (in dark grey) and in the 3-AFC task (in red). Shaded areas represent the standard error to the mean. B – left panel: Average threshold for both tasks, calculated as the mean for the 11 frequencies tested in the 3-AFC paradigm (all vertical lines in panel A). Individual threshold estimates are plotted as dark circles. right panel: Pure Tone Average (PTA) for both tasks, calculated as the mean threshold at 500, 1000, 2000 and 4000 Hz (solid vertical lines in panel A). Individual PTAs are plotted as dark circles. *** indicate significance at p < 0.001*

### Experiment 2

#### Predictive structures induce increased sensitivity

We then tested for what role predictive sequential structure plays in this difference using the fast- and slow-paced paradigms described in the Methods section. In Figure 4A, we first compared the most random version of each paradigm to assess how changes in protocol for each compare across the tasks. A one-way ANOVA shows a significant difference across paradigms (F(3, 81) = 191.91, p < 0.001). Pairwise comparisons reveal significant differences between all paradigms (3AFC vs Randomized: t(27) = −19.55, p < 0.001; 3-AFC vs fast-paced: t(27) = −12.79 p < 0.001; 3-AFC vs slow-paced: t(27) = −17.65, p < 0.001; Randomized vs fast-paced: t(27) = 6.65, p < 0.001; Randomized vs slow-paced: t(27) = 3.68, p = 0.001; fast-paced vs slow-paced: t(27) = −3.02, p = 0.006). The effect of sweeping paradigms compared to the Randomized paradigm reveal that the change in protocols alone - even when cues are reduced as much as possible - provides added information to the presence of the tone for the listener to exploit.

**Figure 4:**
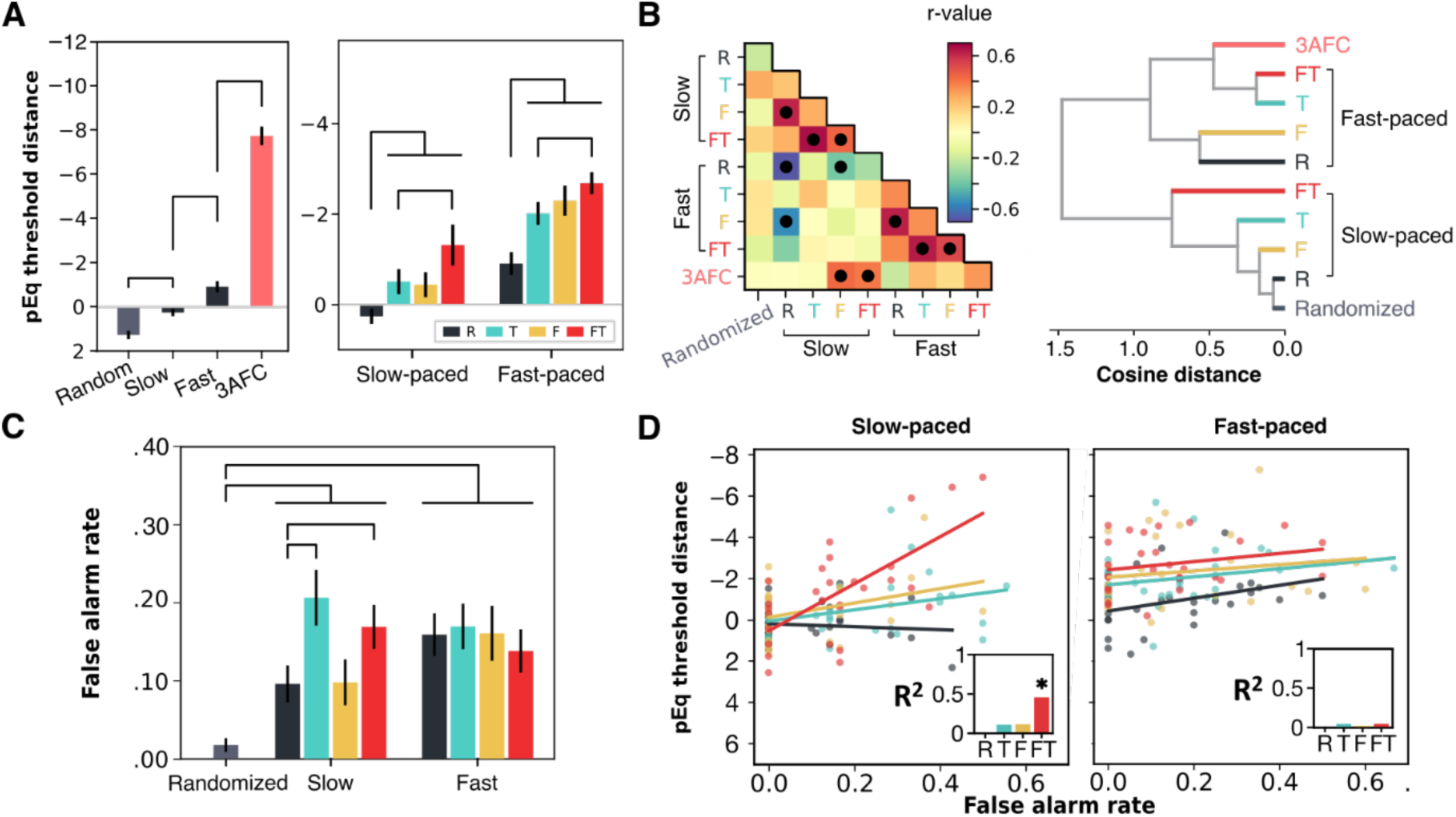
Effect of prediction on detection thresholds. *A – threshold distance effects due to protocol (left) and to predictability (right). Protocol distance is assessed by comparing the random conditions within each protocol. Significant post-hoc comparisons indicated with black lines. B – protocol clustering based on performance correlations across subjects. Left, Correlation matrix of subject performance by protocol conditions. Filled circles indicate significant correlations. Right, Hierarchical clustering of protocols by cosine distance of subject performance in each cluster. C – False alarm rates in each condition. Significant post-hoc pairs indicated with black lines. D – correlation between threshold distance and false alarm rates for continuous (left) and cluster (right) paradigms. Scatter plots indicate single subject plots, lines indicate linear fit between the two parameters, insets reveal R*^2^ *of the correlation between false alarms and threshold distance. * indicate significant relationship*.

Next, we compared the predictable conditions within the fast- and slow-paced paradigms (Figure 4A, right) to see what role prediction plays in auditory sensitivity outside of the protocol differences analyzed above. To this end, we conducted a separate one-way ANOVA for each paradigm, with predictability (Random (R), Timing (T), Frequency (F), or Frequency and Timing (FT)) as the independent variables. For both paradigms, the ANOVA revealed a significant main effect of predictability (Slow-paced: F(3, 81) = 6.92, p < 0.001; fast-paced: F(3, 81) = 14.62, p < 0.001), confirming that even in a tightly controlled paradigm, predictive structure enhances pure-tone detection sensitivity. Post-hoc paired t-tests further showed that all predictable conditions improved performance compared to the most random condition, in both tasks (Slow-paced - FT vs. R: t(27) = −3.60, p = 0.008; F vs. R: t(27) = −3.22, p < 0.001; T vs. R: t(27) = −2.63, p = 0.028; Fast-paced - FT vs. R: t(27) = −6,29, p < 0.001; F vs. R: t(27) = −5.28, p < 0.001; T vs. R: t(27) = −3.85, p < 0.001). Thresholds also improved significantly in the fully predictable (FT) condition compared to the condition with predictable timing only (Slow-paced - FT vs. T: t(27) = −2.39, p = 0.036; Fast-paced - FT vs. T: t(27) = −3.16, p = 0.006). In the Slow-paced paradigm, there was a trend towards significance when comparing the fully predictable (FT) condition to the condition with predictable frequency only (F), although this did not reach significance after correcting for multiple comparisons (t(27) = −2.12, p = 0.052). All other comparisons were not significant (p > 0.234).

We then compared the gain from having both types of predictability together (FT - R) to the combined gains from each type of predictability independently ((F - R) + (T - R)). This comparison allowed us to assess whether the combined predictability provides a greater benefit than the sum of its parts, or whether the effects of frequency and time are independent of one another. In the fast-paced condition, a paired t-test revealed that the combined gain from frequency and time independently was significantly greater than the gain from having both jointly predictable, (t(27) = −2.47, p = 0.020). Conversely, in the slow-paced condition, no significant difference between these gains was found, (t(27) = 0.32, p = 0.749). These results suggest that in the slow-paced task, the gain from predictability in each dimension is likely independent. However, in the fast-paced task, the gain from the sum of both predictabilities independently is larger than the gain from having both frequency and time jointly predictable, hinting at a different integration mechanism between the two tasks.

#### Distinct predictive sources across timescales

We next investigated how performance correlated between task pairs. Our aim was to test whether performance was grouped by the category of predictability (frequency vs timing) or by the paradigmatic setup (slow- vs fast-paced). The correlation matrix, shown in Figure 4B, reveals that the structure of the task most drives similar performance rather than predictability in a particular dimension like time or frequency. This suggests that the two different time scales lead to different mechanisms through which predictive structure can support the analysis. To test this further, we included participant performance into an unsupervised clustering algorithm (4B, right) to assess whether the patterns of participant performance could allow inferring similar or different mechanisms across paradigms. The clustering algorithm clearly grouped performance by paradigm rather than by the type of predictability in the condition.

#### Predictive gain mostly driven by sensitivity, not decisional bias

Another important question is what role decisional bias may play in this increase in performance. As the Experiment 2 paradigms involved a detection task, it could be that participants claimed detection more even when they heard nothing. To test for this, we included catch trials in the experiment during which no tone occurred. Figure 4C reveals the results of the catch trial analysis. We conducted a repeated measures ANOVA with the false alarm rate as the dependent variable, treating each condition as a separate group to allow for comparisons among all conditions. The analysis revealed a significant effect of group (F(8, 216) = 5.78, p < 0.001). Post-hoc paired t-tests using FDR correction indicated that false alarm rates in the Randomized paradigm were significantly lower than in all conditions of Experiment 2 (Randomized vs Fast F: t(27) = −3.84, p = 0.004; Randomized vs Fast FT: t(27) = −4.21, p = 0.002; Randomized vs Fast R: t(27) = −5.00, p < 0.001; Randomized vs Fast T: t(27) = −5.02, p < 0.001; Randomized vs Slow F: t(27) = −2.58, p = 0.047; Randomized vs Slow FT: t(27) = −5.13, p < 0.001; Randomized vs Slow R: t(27) = −3.15, p = 0.018; Randomized vs Slow T: t(27) = −5.56, p < 0.001). Within the slow paradigm, false alarm rates were higher in both conditions of the task with predictable timing compared to the random condition (FT: t(27) = 2.97, p = 0.022; T: t(27) = 3.22, p = 0.017), and they were lower when only the frequency was predictable than when only the time was predictable (F vs. T: t(27) = −3.07, p = 0.019). While temporal predictability in the slow paradigm does appear to be modulated by decisional bias, it is unlikely to explain the entirety of the result. Predictability in frequency does not appear to affect false alarms (F vs R: t(27) = −0.15, p = 0.965) while still improving detection thresholds. At the same time the fast paradigm shows a nonsignificant reduction in false alarms as predictability increases (F(3, 69) = 0.668, p = 0.575), excluding decisional bias as a possible explanation of the result. To further test this hypothesis, we compared false alarm rates of individual participants against their threshold differences. The slow-paced paradigm shows a significant correlation between false alarm rate and threshold detection when time and frequency are predictable, indicating some contribution of decisional bias to threshold detection (FT: R^2^ = 0.456, p < 0.001). However, the other conditions showed no significant correlations (R: R² = 0.01, p = 0.604; T: R² = 0.109, p = 0.086; F: R² = 0.116, p = 0.077). There is also no such significant correlation in the fast-paced paradigm (R: R² = 0.116, p = 0.076; T: R² = 0.051, p = 0.247; F: R² = 0.026, p = 0.408; FT: R² = 0.052, p = 0.244).

#### Individual differences not explained by musical experience

We then sought to explain the variability in participants’ ability to extract predictive information in terms of other characteristic features that we had tracked through surveys: musical expertise and age (Fig. 5). Neither feature had a particularly strong effect on participant performance. In the continuous paradigm, Age had no effect on threshold distance (R: R² = 0.001, p = 0.878; T : R² = 0.001, p = 0.872; F : R² = 0.008, p = 0.658; FT: R² = 0.000, p = 0.914). In the cluster paradigm, Age had a worsening effect on Random and Time conditions but no effect on Frequency and Both conditions (R: R² = 0.154, p = 0.039; T: R² = 0.226, p = 0.011; F : R² = 0.007, p = 0.68; FT: R² = 0.011, p = 0.601). Surprisingly, Musicianship (as indexed by the General Sophistication score of the GMSI) had no effect on threshold distance in any condition (Fast – R: R² = 0.017, p = 0.508; T: R² = 0.001, p = 0.861; F: R² = 0.061, p = 0.204; FT: R² = 0.008, p = 0.653; Slow – R: R² = 0.026, p = 0.416; T: R² = 0.002, p = 0.827; F: R² = 0.023, p = 0.438; FT: R² = 0.003, p = 0.776).

**Figure 5:**
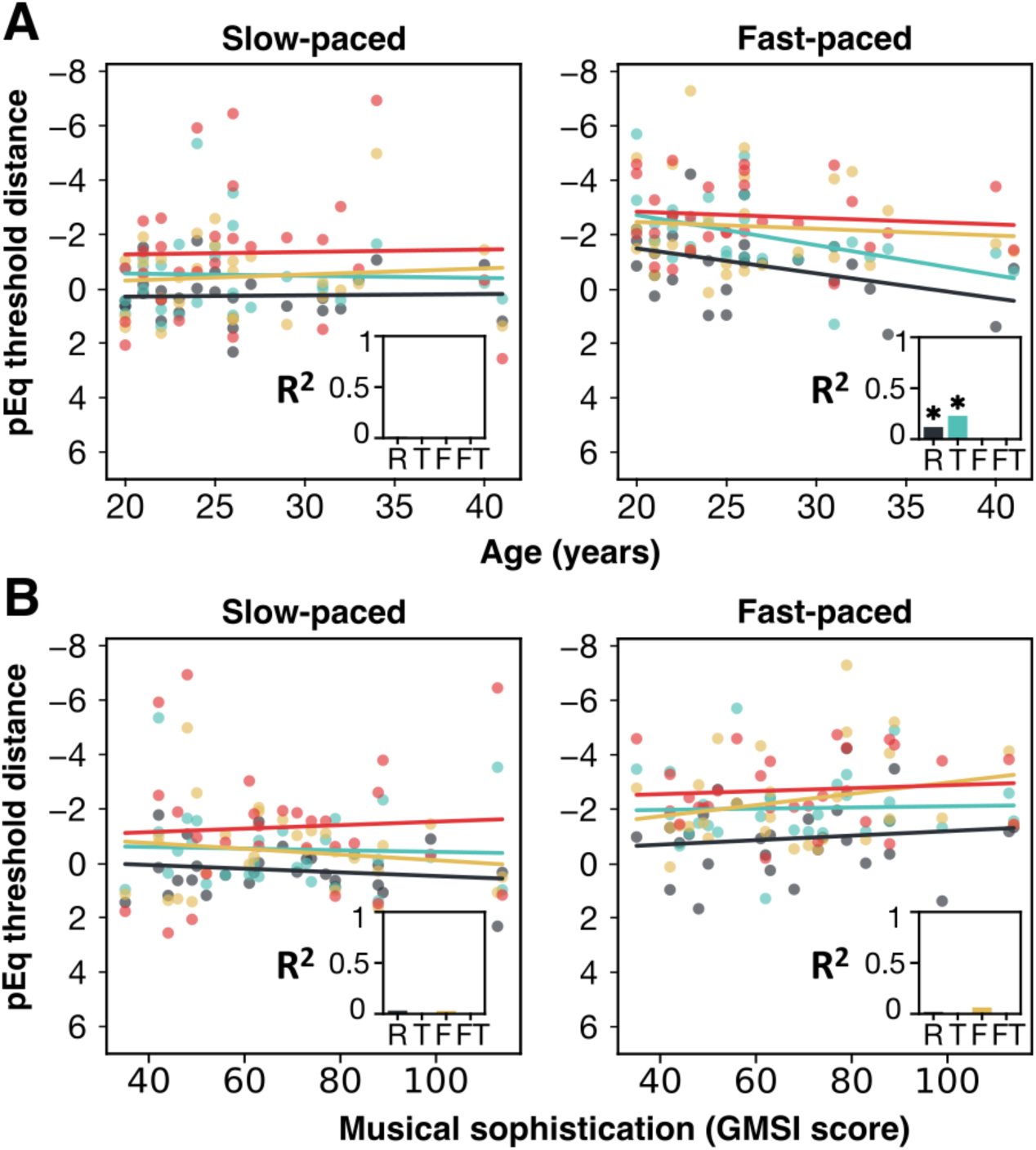
Predictive effects variability largely unexplained by age and musicianship. *A – correlation between threshold distance and participant Age, in continuous paradigm. Scatter plots indicate individual participants, lines indicate linear regression. R*^2^ *of variance explained is reported in upper right insets. Asterisks indicate significance. B – Correlation between threshold distance and participant musicianship. Plot organization is the same as in A*.

## Discussion

Our findings suggest that the human brain uses predictive information to enhance pure tone auditory sensitivity even at detection threshold, previously thought to be a largely low-level peripheral process. In experiment 1, our least and most predictable tasks (the Randomized to the clinical 3AFC paradigm), we found a 7-dB average difference within the same subjects. This effect is sizeable when compared against the 5-dB step size used for clinical measurement (32). We then used more high-resolution methods to identify how predictive structure in time and frequency contribute to this difference. We found evidence that predictability in both frequency and time contribute substantially to this predictive effect at two timescales, a slow pace consistent with audiological protocols (1 - 3 seconds per tone) and a faster pace consistent with time scales of sequences of acoustic events like speech and music (400 - 800 ms). Our findings reveal a contextual influence on peripheral *sensitivity*, likely instantiated via top-down modulation by feed-back connections, an effect that remains insufficiently explored and assessed in audiological fields.

Our study investigated the role of prediction under two scenarios: a slow-paced condition in which participants continuously report detection on every tone they hear, spaced out about 1-3 seconds, and a faster-paced condition, whereby participants first hear a cluster of cue tones (400-800 ms between them) which cue predictively (or not) the timing or frequency (or both) of a target detection tone. Our strongest results occur in the fast-paced case. This finding is in keeping with the previous literature on prediction. In this faster paced case, tones are detected as part of a sequence occurring at a pace typical of music (∼1.5-3 Hz). Previous work has shown that predictions are strong within these sorts of sequences and peak particularly at around this time scale (18, 33). While the slow-paced task also consists of a sequence of tones, the amount of time spaced between tones is quite large making it more likely to be processed as isolated events. Previous work has shown that particularly in non-musicians, sequences are processed differently when item rate is less than 1 Hz (34).

In our results, we observed that in the slow-paced paradigm, the gain from having both – frequency and time– predictabilities independently was not significantly different from having the two together, suggesting the two forms of prediction linearly combine. This implies that at a slower pace, participants are likely processing these predictabilities independently, perhaps due to the longer intervals allowing for more isolated processing of each predictability cue. Conversely, in the fast-paced paradigm, the gain from the sum of both predictabilities independently was significantly greater than the gain from having both frequency and time jointly predictable. This suggests that at faster paces, the predictabilities interact in a redundant manner when they are combined, hinting at potentially distinct underlying mechanisms. The faster-paced task may, therefore, engage a more complex integrative mechanism, possibly due to the temporal dynamics aligning more closely with naturalistic auditory processing such as in music.

Further proving that the two protocols reveal distinct mechanisms of prediction is our clustering analysis. We initially expected that, regardless of pacing, participants would be grouped by the type of prediction (frequency or time). Instead, we found the opposite effect, participant performance was best clustered by the time-scale. Furthermore, each predictable condition within each protocol clustered together, away from the random condition. These findings suggest that similarity across participants is driven more so by prediction ability at distinct timescales rather than over distinct features. This clustering is not due to the overall mean shift in performance across the protocols. Our clustering method operates over the cosine similarity across participants, which ignores this overall shift.

Our results exclude the possibility of two alternative hypotheses to explain the improvement in thresholds: 1) a change in decisional biases and 2) a change in cochlear state. A change in decisional bias would plausibly lead to improved thresholds if for example participants indiscriminately indicated detecting the sound regardless of the stimulus. We address this concern by including catch trials in which no sound was presented. By analyzing these trials, we find that decisional bias cannot explain our results. While in the slower paced experiment, false alarms increased with temporal prediction, predictability in frequency shows no increase in false alarms. In the faster-paced experiment, false alarms go down with predictability (though not significantly). From this we can negate the first alternative hypothesis: a significant portion of our results must be due to an increase in *sensitivity* not to a change in decisional bias.

The second alternative hypothesis suggests that the increase in sensitivity is due not to central processing but instead to a local change in the cochlear processor. A myriad of studies have shown that local peripheral responses can change as a result of preceding and concurrent stimuli including masking effects (35), and noise protection through the modulatory effect of the Medial Olivocochlear Reflex (36). Our experimental design also negates this possibility as the gain in performance is demonstrated in comparison against a random control which maintains the stimulus context in every way except predictability. For example, comparing the effect of predictability in timing vs random, in this case the frequency distributions of tones, the temporal intervals between tones and the visual cue for the target window are exactly the same. The only change between these two conditions is that the temporal intervals are held constant per trial in the predictable case and shuffled for the random. To take advantage of this, the cochlea would need to execute high-level computations that are typically expected in cerebellum, basal ganglia and motor cortex (for a recent review, see 37). From our perspective, a temporal prediction redundancy in the cochlea is a far less plausible hypothesis.

Instead, we propose that our findings result from an interaction of top-down predictive information with cochlear responses, enhancing sensitivity for specific frequencies and time points based on expectation. Whether this interaction involves a change in cochlear processing directly or instead of the central interpretation of cochlear responses will require future work. However, either hypothesis has major implications for the fields of audiology and neuroscience. For example, if the hypothesis of altering cochlear responses via central input is correct, it would provide a mechanism for phase locked loops whereby the central perceptual system can endogenously and voluntarily alter its bottom-up input, similar to eye movements or whisking but in the spectral domain (38–40). Alternatively, if the results are due to a change in central interpretation, the findings upend the assumption in audiology that pure tone audiogram measure purely cochlear health (13).

While audiologists are aware that predictability can alter their results, – it is standard practice in several countries to present tones with random timing, for example – this effect is thought to be due to our first alternative hypothesis: decisional bias, patients may report detecting a sound regardless of what they have heard, purely because they know where the tone should have been. Here we show for the first time that prediction can not only alter decisional bias but also enhance sensitivity and to a significant degree. Therefore, depending on the chosen clinical set up of the paradigm, the pure tone audiogram may be diagnosing more fully the auditory neural apparatus, mixing effects of both peripheral cochlear health and central, predictive ability. From the perspective of standard practice, this finding may not represent a significant issue for clinicians: the typical sounds that patients experience in their lives have some degree of predictability in them and as such current clinical setups match patients’ daily experience. Still, our findings reveal two separable components present in sensory detection which may have separate trajectories of development and degradation over the course of lifespan. Future work will investigate these trajectories more fully.

To that end, while the current task is not designed to study the individual differences in perception, we used the natural variance in our participant cohort to assess how age (<45 yo) could affect the predictive trajectory. The only significant effect we found was in the random and timing predictions of our fast-paced task. Interestingly predictability in frequency (F and FT) removed this effect, suggesting that even in younger populations, age reduced sensitivity but that this effect could be masked by improved predictability. Future work will assess in more detail the trajectory of this effect both with greater age and with clinical sensory hearing loss to assess how prediction can be used either as a coping mechanism or an indicator of future decline.

Together, our findings reveal that pure tone audiometric measurements contain the influence of both bottom-up sensory responsivity and top-down predictive modulation. We show that prediction in both frequency and time and at multiple timescales can influence the sensitivity of the auditory system. Signal detection is most improved in the context of sequences with timing similar to those of typical auditory stimuli like speech and music. We also show key differences in behavior between these timescales including that temporal predictions at slower timescales may influence more decisional bias whereas frequency predictions improve sensitivity. Lastly, we look for individual differences to understand the variability in predictive performance. Even in our younger population, we find an effect of age on sensory detection in fast paced sequences which can be recovered by frequency predictability. We expect our findings to inspire audiologists and auditory neuroscientists to track raw predictive ability more carefully across age and lifespan as a potential key factor in understanding hearing abilities.

## Author contributions

NM, DSL, LHA and KBD designed research. NM and KBD performed research and analyzed data. NM, LHA and KBD wrote the first draft of the paper. NM, GG, HJ, NP, NW, DSL, LHA, KBD edited the paper.

## Acknowledgements

The authors would like to thank Benjamin Morillon, Céline Quinsac, Paul Avan, and Lavinia Slabu for their insights and discussions on this project.

This work was supported by the Fondation pour l’Audition (RD-2020-10, LHA), a French government grant managed by the Agence Nationale de la Recherche under the France 2030 program (ANR-23-IAHU-0003, LHA), a Fondation Fyssen Postdoctoral Fellowship (KBD) and BPIFrance i-Nov grant DOS0127610/00 (NW).

## Supplemental information

Document S1. Details on the iAudiogram procedure

## Resource Availability

### Lead contact

Requests for further information and resources should be directed to and will be fulfilled by the lead contact, Keith B. Doelling.

### Materials availability

This study did not generate new unique reagents.

### Data and code availability

The anonymized data and the code for analysis have been deposited in a public GitHub repository and are available at https://github.com/pimpimpula/ThereWillBeBeeps.

## Supplemental Materials

### S1: Details on the iAudiogram procedure

The procedure starts by collecting initialization data, testing tones spanning the frequency range (1000 Hz, 1500 Hz, 2000 Hz, 3000 Hz, 4000 Hz, 6000 Hz, 8000 Hz, 750 Hz, 500 Hz, 250 Hz, 125 Hz) to provide an initial coarse estimation of the audiogram. After each tone presentation, the algorithm requires a positive or negative answer to select the next stimulus. During the initialization phase, the frequency of the tones is predictable, as one frequency is tested several times in a row at decreasing hearing levels, until a negative answer is recorded for that frequency. To avoid repeating this phase in the two Sweeping paradigms where we aim to control the predictability of oncoming tones, we reused the initialization data recorded during the Randomized task to start all the audiograms measured in the Sweeping paradigms.

